# Vibrational noise pollution in bee hives generated by railway traffic

**DOI:** 10.1101/2025.07.15.664893

**Authors:** Sarah Chehaimi, Marie Christine Seidel, Jennifer Richter, Wolfgang H. Kirchner

## Abstract

Anthropogenic noise pollution has become a threat for the fauna with possible effects on animal physiology and behaviour. We explored the effects of anthropogenic vibrational noise on honey bees (*Apis mellifera*). The intensity of substrate borne vibrations generated by trains on the ground, on the front comb of a hive and on its base as well as the airborne sound pressure were recorded at different distances and sites. Vibrational noise of amplitudes that can be detected by honey bees is present on the ground surface, on the comb and on the base over distances up to 20 m from the railway track and the attenuation of the airborne sounds is higher than the attenuation of the substrate borne vibrations. Bees placed in an observational hive and exposed to simulated substrate borne vibrations caused by a cargo train show significant behavioural reactions. The same substrate borne vibration was presented to bee colonies every 5 minutes continuously for six months at realistic amplitudes. On the colony level no differences are found for capped and open worker brood, worker eggs, adult drone population, capped and open drone brood, brood development, varroa mites, collected pollen and honey production; while some significant differences are present for worker population and pollen collecting foragers. We conclude that honey bee colonies in the vicinity of railroads are exposed to substrate borne vibrational noise above their threshold of sensitivity up to 20 m. At the individual level bees show reactions to the vibrations; however, at the population level bees seem to cope with the disturbance. As higher amplitudes and additional stress factors might affect the colonies, it seems to be anyways advisable to generally avoid placing bee colonies close to anthropogenic vibrational noise sources.

## 1. INTRODUCTION

The anthropogenic colonization of terrestrial and aquatic environments has contributed enormously to the global pollution. Anthropogenic mechanical noise pollution is one of its by-products and is intended as any external vibro-acoustical stimulus generated by human activities, which could limit signal or cue detection (Roberts&Howard 2022) and/or have destructive consequences on humans and animals (Slabbekoorn 2019). The anthropogenic mechanical noise differs consistently from the natural noise of rain, wind, conspecifics or heterospecifics (Raboin&Elias 2019); it can be impulsive or continuous, mobile or stationary and may vary in frequency, amplitude and occurrence (Roberts&Howard 2022), often overlapping the natural frequencies (Virant-Doberlet et al. 2014). Moreover, it can travel throughout various media (air, water, substrates) and can be conveyed from a medium to another. The negative impact of mechanical noise pollution has been reported by a growing number of studies (European parliament 2002; Rabanal et al. 2010; WHO 2011; Ortega 2012; Williams et al. 2014; see review Classen-Rodriguez et al. 2021; see review Roberts&Howard 2022); however, little is known specifically about the effects of vibrational noise pollution on invertebrates. Insects use substrate borne vibrations for communication alone or in combination with other forms of mechanical signalling (Cocroft&Rodriguez 2005). The frequencies most used by insects are frequencies below 1000 Hz, which are subjected to lower attenuation (Bennet-Clark 1998; Čokl&Virant-Doberlet 2003) but highly overlapping with the frequency range of anthropogenic vibrational noise (e.g. airport traffic and railways are known to produce intense vibrations below 1000 Hz; Ono&Yamada 1989; Heckl et al. 1996; Fidell et al. 2002). The co-occurrence of communicational signals and anthropogenic mechanical noise can lead to masking, distraction, misleading and/or development of strategies by animals to cope with the disturbance, like competition for vibro-acoustic space, plasticity for frequency and intensity level (when possible as insects are limited due to their own body characteristics) or negative impact on the youths (for review, see Classen-Rodriguez et al. 2021).

In our knowledge, vibrational noise pollution in relation to honey bees (*Apis mellifera*) has never been explored. They have evolved diverse vibrational signals in order to efficiently communicate within the colony (Allen 1956; Wenner 1962; Simpson 1964; Seeley 1985; Michelsen et al 1986b; Kirchner 1993; Nieh 1993; Ramsey et al. 2018; Hrncir et al. 2019; see review Kirchner et al. 2022), therefore they may be target of vibrational disturbances, especially in Germany where they are often found nearby railroads. The subgenual organ, located in the tibia of the legs, which detects the substrate borne vibrations, is particularly sensitive to the dorsoventral component of the vibration (Kilpinen&Storm 1997, Rohrsitz&Kilpinen 1997, Rohrseitz 1998). Electrophysiologically, the lowest vibrational threshold is to be about 0.1 m/s² at 300 Hz (0.06–0.15 mm/s peak-peak, Kilpinen&Storm 1997). Behaviourally, the lowest vibrational threshold for eliciting a freezing response in a hive is found to be about 4 m/s² at 200 Hz (2 mm/s peak-peak; Rohrsitz&Kilpinen 1997) and about 5 m/s² at 300 Hz (3 mm/s peak-peak; Michelsen et al. 1986a). In a recent study (Chehaimi&Kirchner 2024), the vibrational threshold measured behaviourally in an isolated arena has been found as low as 0.25 m/s² in virgin queens and workers, 0.55 m/s² in laying queens and 0.6 m/s² in drones at 500 Hz and, respectively, about 0.3, 0.4, 1 and 1.8 m/s² at 350 Hz. The vibro-acoustic waves detected by bees inside the hive are modified by the hive itself, which functions as a resonator (Rayleigh 1916), and its inner soundscape (collection of all detected and interpreted sounds by an organism; Farina et al 2021) is a composition of bee buzzing (sounds generated by voluntary and involuntary beeś behaviour) and external sounds (Farina 2023). There is increasing evidence that honey bees are able to use the combination of external sounds and inner buzz as an informative medium for communication and coordination of social activities (Terenzi et al. 2020; Farina 2023) and to extract ecoacoustic codes (set of codes used to describe sounds and vocalization of other animals in their natural environment; Farina&Villa 2023). This opens the possibility that bees may cope with an external disturbance till a certain extent and distinguish their own vibro-acoustics from external ones. Few experiments (Frings&Little 1957; Little 1962; Stefanec et al 2021) showed that honey bees react differently to the disturbance (freezing, slowed activity performance after initial freezing, increased activity, diminished activity) depending on its frequency. Other experiments showed effects of vibrational disturbance on the colony level in terms of flight reduction (up to 80%; Spangler 1969) and delay of queen emergence (Spangler 1971). Bencsick et al. (2024) used the delivery of a short (0.1 s) vibrational disturbance to assess the physiological status of the colony (e.g. queenless versus queenright colony) based on the quantification of its mobility: during the pulse, the bees are immobilized and after the pulse, the bees recover exponentially.

We therefore started to investigate whether the railway traffic generates substrate borne vibrations on the ground and on a hive at intensities that can be detected by honey bees and whether the vibrational noise has an effect on individual bees or honey bee colonies.

## 2. MATERIALS AND METHODS

### Recording of sounds and vibrations close to railroads

The experiment took place between winter and early spring in 2021-2022 and in winter 2022-2023. Four measuring sites have been selected in the German federal state of North Rhine-Westphalia: Bochum (54% cargo trains, 30% regional trains, 16% locomotives), Düsseldorf A (67% ICE, 30% regionals, 3% locomotives), Düsseldorf B (83% regionals, 11% local trains S-Bahn, 6% locomotives), Dinslaken (39% cargos, 38% regionals, 9% ICE, 9% S-Bahn, 5% locomotives). The sites differed in landscape, ground type and level of the railway. The ground in Bochum, Düsseldorf A and Düsseldorf B was made of soil and was characterized by the presence of bushes and/or trees after 10 or 20 m. In Dinslaken the ground was made of stones and the environment was free from obstacles. The railway was minimally raised in Bochum and Düsseldorf A at 5 m, while it was even in Düsseldorf B (ascendant only at 10 m) and Dinslaken. The substrate borne vibrations generated by the passage of trains were recorded in all sites, while the airborne sounds were recorded only in Düsseldorf A, Düsseldorf B and Dinslaken. The measurements were collected between 9 am and 4 pm, when the railway traffic was more intense. The substrate borne vibrations travelling on the ground were recorded at 5, 10, 20 m distance from the railway track using the accelerometer type KD37 (MMF), which was eased on the ground surface without any support or gluing materials. An empty transportable bee hive (52,5×32,5×30 cm) was then placed in direct contact with the ground and parallel to the railway at 5, 10, 20 m and a KS943B10 tri-axial accelerometer (MMF) was inserted in the middle of the front comb (railway side) to measure the substrate borne vibrations throughout the comb in three dimensions (X=horizontal axis parallel to the comb plane, Y=vertical axis parallel to the comb plane, Z=perpendicular axis to the comb plane). Additionally, a KD37 accelerometer (MMF) was placed on the front internal base protruding toward the inside (52,5×9×2 cm) of the hive with the help of bee wax. The accelerometers were connected via IEPE100 amplifiers (MMF) and SA-P48/CCP-C signal converters to a H6 Recorder (Zoom Corporation). Two microphones MM1 (Beyerdynamic) were positioned at 5 and 10 m, while at 20 m we measured the airborne sounds with a SL10 sound level meter (Voltcraft). The microphones were connected to a DR-40 Linear PCM recorded (Tascam). The intensities (peak-peak) of the recorded vibrations were obtained using the software Audacity 2.4.2. and the substrate borne and airborne amplitude spectra using Raven Pro 1.6. The statistics were conducted with Matlab using U-test (for independent and non-parametric data).

### Playback experiments

An observational hive was setup in the Botanical Garden of Ruhr University of Bochum. A recording of a cargo train (from Dinslaken) was presented to the bees randomly alternated to a control (absence of the vibration) every 5 minutes at 3 m/s² (N trials=44; Chi-square-four-field test), in order to note any behavioural reaction at the individual level during and after the train passage. The recording had 23 s duration and was presented using an audio player MM-2218 (Bässgen) and played back using a CS-PA1MK amplifier and a Visaton vibrational exciter. The experiment was divided in two parts: a pause of 1 h was inserted after 27 trials to avoid possible effects due to the continuous presentation of the vibration (i.e. habituation), despite the trial interval. A baseline behaviour was noted before the first part of the experiment was started and also after the pause before the second part of the experiment was started. In this way any behavioural change in respect to the baseline was considered a reaction. The first baseline behaviours, observed visually (eyes) and acoustically (headphones) by the experimenter, included fanning with the wings, smoothly walking around, resting on the glass, stretching the legs, grooming, antennal interaction, trophallaxis, putting head in the cell and buzzing. In the second baseline assessment, the behaviours observed were the same, but bees were more excited and more engaged in activities as expected in the very first afternoon and when the weather is optimal. The behavioural reaction was evaluated as an increase or decrease in activity and movement in respect to the baseline noted and was again visually and acoustically subjectively evaluated by the experimenter. The area of observation was a circle of 63,6 cm² and was confining about 50 bees. The substrate borne vibrations reaching the area were previously measured with an accelerometer to assure that they had effectively the desired amplitude (3 m/s²).

During spring and summer 2023, 10 colonies (C1-C5 control: ordinary bee colonies; V1-V5 experimental: ordinary bee colonies but subjected to substrate borne vibrations) have been located at the Botanical Garden of Ruhr University of Bochum. The hives were eased down on europallets, sustained by parallelepiped-shaped stones, to insulate them from the ground. The colonies were carefully prepared, so that the two groups had similar characteristics in terms of population (adult workers, capped worker cells, worker larvae and eggs, adult drones, capped drone cells, drone larvae and eggs, honey and pollen stores). The recording of the cargo train (duration 23 s) used for the individual level experiment was also chosen for this experiment as a playback. The recording was looped continuously every 5 minutes using an audio player MM-2218 (Bässgen). The outputs of the audio-player were connected to CS-PA1MK amplifiers and to vibrational exciters (Visaton), placed at the top of each hive onto the wooden lids, from which the vibrations could spread all around the hive. KD37 accelerometers (Metra Meß-und Frequenztechnik), connected to MMF IEPE100 preamplifiers, were inserted in the central comb of each hive and connected to an oscilloscope to monitor the vibrational amplitudes.

When the season was proceeding and other honey boxes were added, this comb was always situated in the box just above the queen excluder. The amplitude was initially set at 1 m/s² (from 20^th^ March to 16^th^ July 2023) but successively risen up to 3 m/s² (from 17^th^ July to 22^nd^ September), to see whether more evident effects would occur (see Results section). Hive scales (We Gro) were placed below each hive and automatically recorded the total weight of the hives daily at 9 pm. The colonies were managed following standard beekeeping treatments (cutting of drone frames, swarm control, harvesting honey, feeding for the winter cluster). The population was re-estimated every 3-4 weeks using the Liebefeld method for Zander hives. Following this method, each comb is divided mentally in 8 areas of equal size and each area full of bees contains approximately 125 bees. This reference permits to estimate the bee population located on the combs, to which the bee population standing on the frames and walls must be added (Imdorf et al. 1987). Following the OECD 75 guideline for the assessment of effects of plant protection products on honey bees (OECD 75, 2014), brood development in terms of brood index, termination index and compensation index was also monitored by taking photographs of 100 cells for each of the two sides of one selected comb per colony every 0, 5, 10, 16, 22 days. The brood index indicates the development rate of the brood from the status of egg till the status of empty cell (when the adult bee has emerged or when the cell is replaced with another egg). The termination index indicates the death rate of the larvae and pupae during the development process. The compensation index indicates the capacity of the colony to replace undeveloped larvae and pupae. The acquired photos were analysed using the software HiveAnalyzer v 2.32 (Visionanalytics). Moreover, the number of returning foragers with pollen was also monitored every week at 12 am for 1 minute from April onward. In September, evaporated oxalic acid against *Varroa destructor* was applied to all colonies with intervals of about 5 days. The statistical analysis was run with Matlab using U-test (for independent and non-parametric data) to compare the experimental and control groups.

## 3. RESULTS

### Recording of sounds and vibrations close to railroads

Trains passing bee hives can induce strong substrate borne vibrations throughout the comb with amplitudes up to more than 4 m/s² peak-peak in Y and Z axis at 5 m (Fig. 1A). Most of the energy of train traffic is concentrated in the frequency range below 1 kHz to which bees are most sensitive (Fig. 1B).

**Fig. 1.**
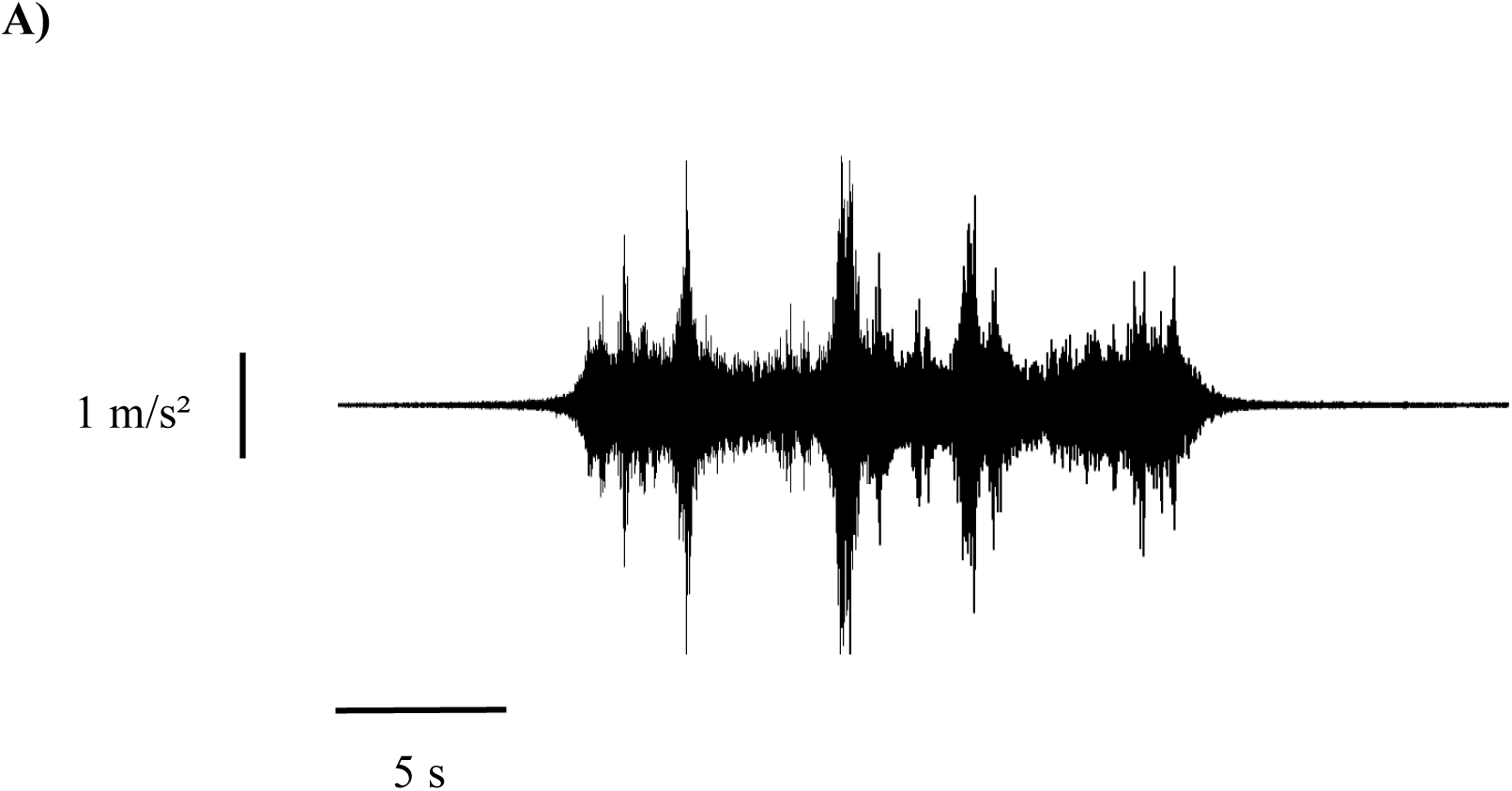
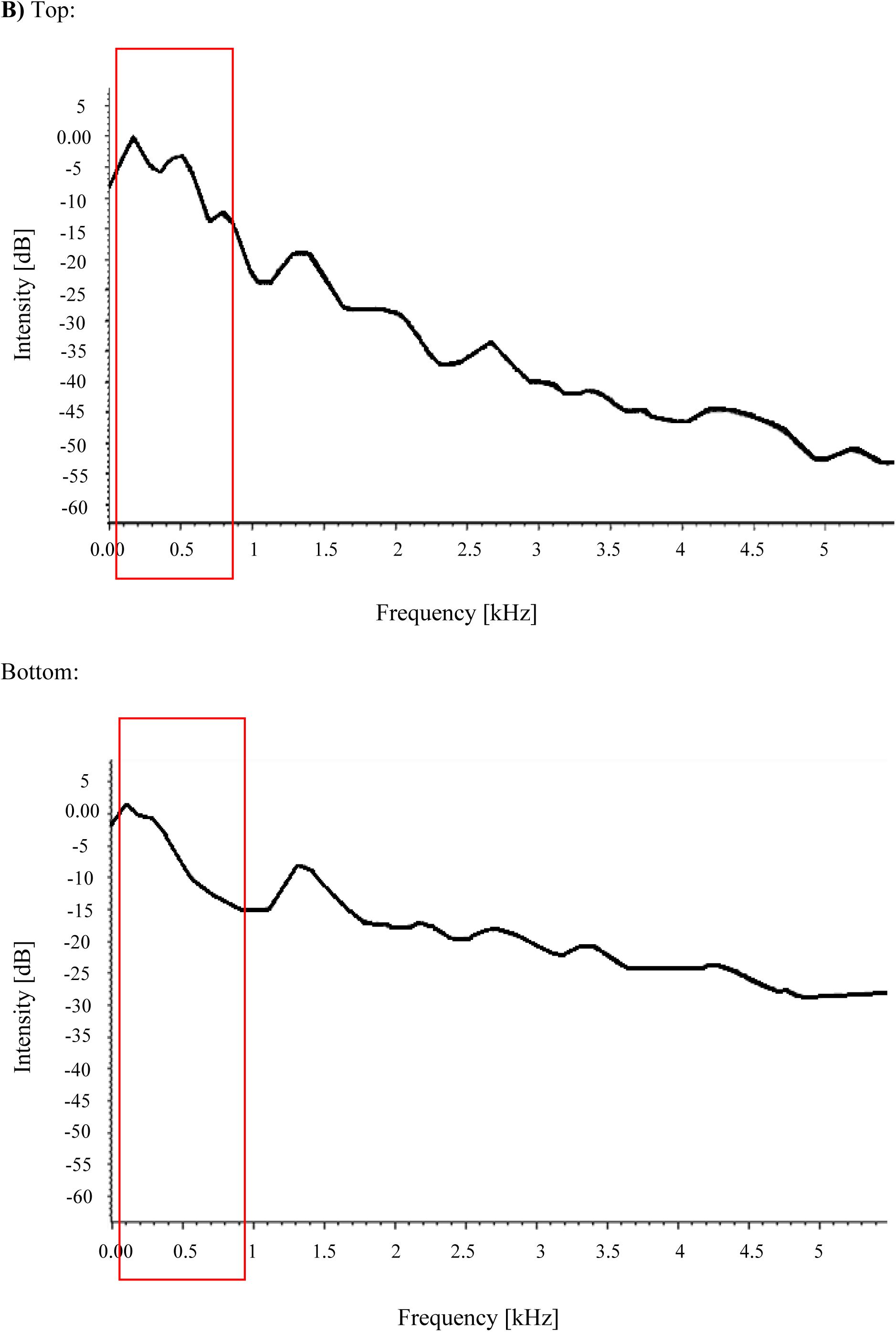
A) Vibrations in Y-axis generated by the passage of a cargo train at 5 m: maximal amplitude 4.4 m/s² peak-peak. B) Top: substrate borne amplitude spectrum. Bottom: airborne amplitude spectrum. Most of the energy of train traffic is concentrated in the frequency range below 1 kHz, where the frequencies used by honey bees for substrate borne vibrational communication fall (red square).

The highest vibrational amplitudes were found in the Z axis on average, perpendicular to the surface of the comb (N trains=182; Fig.2A, B left), which coincides with the directional component most perceived by honey bees (dorsoventral component; Kilpinen&Storm 1997). The intensity of the induced vibrations is above the vibrational sensitivity threshold of honey bees. We compared the substrate borne vibrations induced by trains in the different sites (N trains=78; Fig. 2B center). The highest average intensity (Z axis) was found in Dinslaken (1.9 m/s²) and the lowest in Düsseldorf B (0.8 m/s²) at 5 m. The vibrations measured at the base of the hive are significantly higher than the ones measured on the ground itself (N trains=439; Fig. 2B right).

**Fig. 2.**
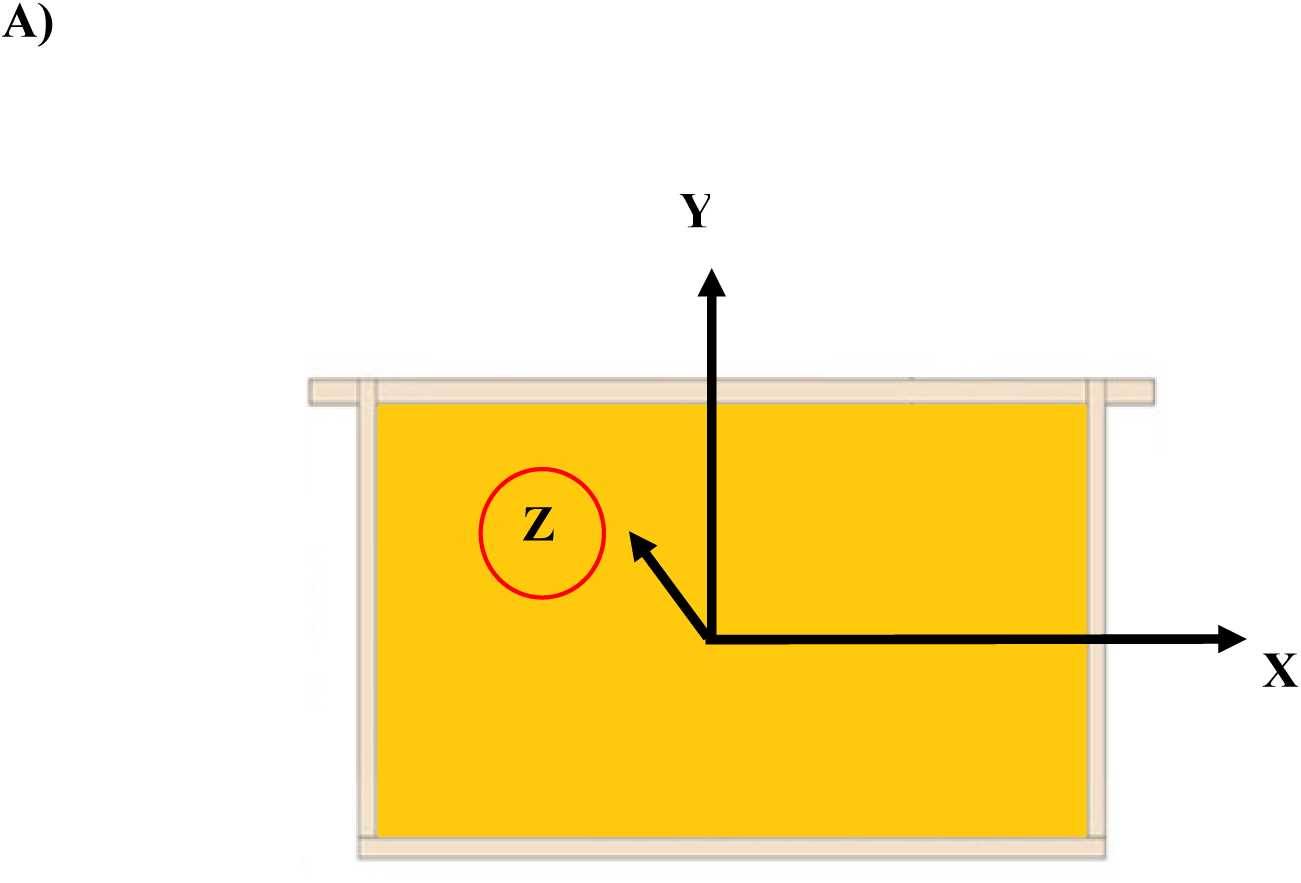
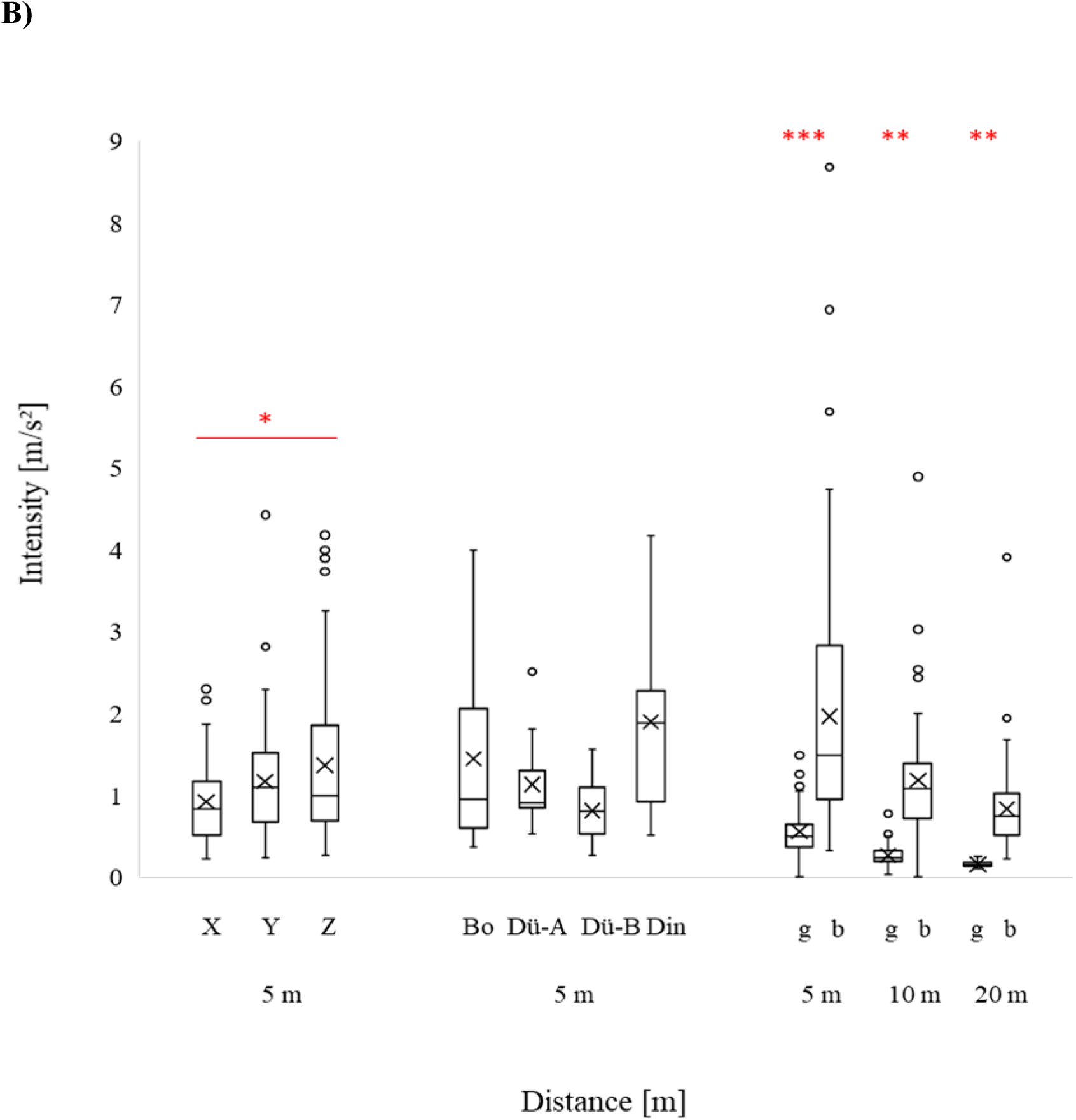
A) Representation of the three axes (X, Y, Z) in respect to the comb. The Z axis, perpendicular to the comb, corresponds to the dorsoventral axis of the bee leg, therefore coinciding with the directional component most perceived by the bees. B) Left: comparison of vibrational amplitude [m/s²] between X, Y, Z axis from all sites at 5 m in the hive: Z is the most vibrated axis on average (N trains=182; U-test; X-Y: p>0.05; X-Z: p<0.05; Y-Z: p>0.05). Center: vibrational amplitude [m/s²] for the Z axis in the different sites at 5 m distance from the railway (N trains=78; Bo=Bochum; Dü-A=Düsseldorf A; Dü-B=Düsseldorf B; Din=Dinslaken). Right: comparison of the average amplitude [m/s²] of ground (g) and hivés base (b) vibrations of Düsseldorf A, Düsseldorf B and Dinslaken (N trains=439; U-test; 5 m: p<0.001; 10 m: p= 0.001; 20 m: p=0.001).

The vibrations extend up to 20 m with an average intensity of 0.7 m/s² (96,9 dB) for all sites, still overlapping the range of vibrational sensitivity of honey bees (N trains=679; Fig. 3). The attenuation of the airborne sounds is 6 dB per doubling of distance plus some variable linear excess attenuation, while the attenuation of the substrate borne vibrations is 3 dB per doubling distance plus a small excess attenuation.

**Fig. 3.**
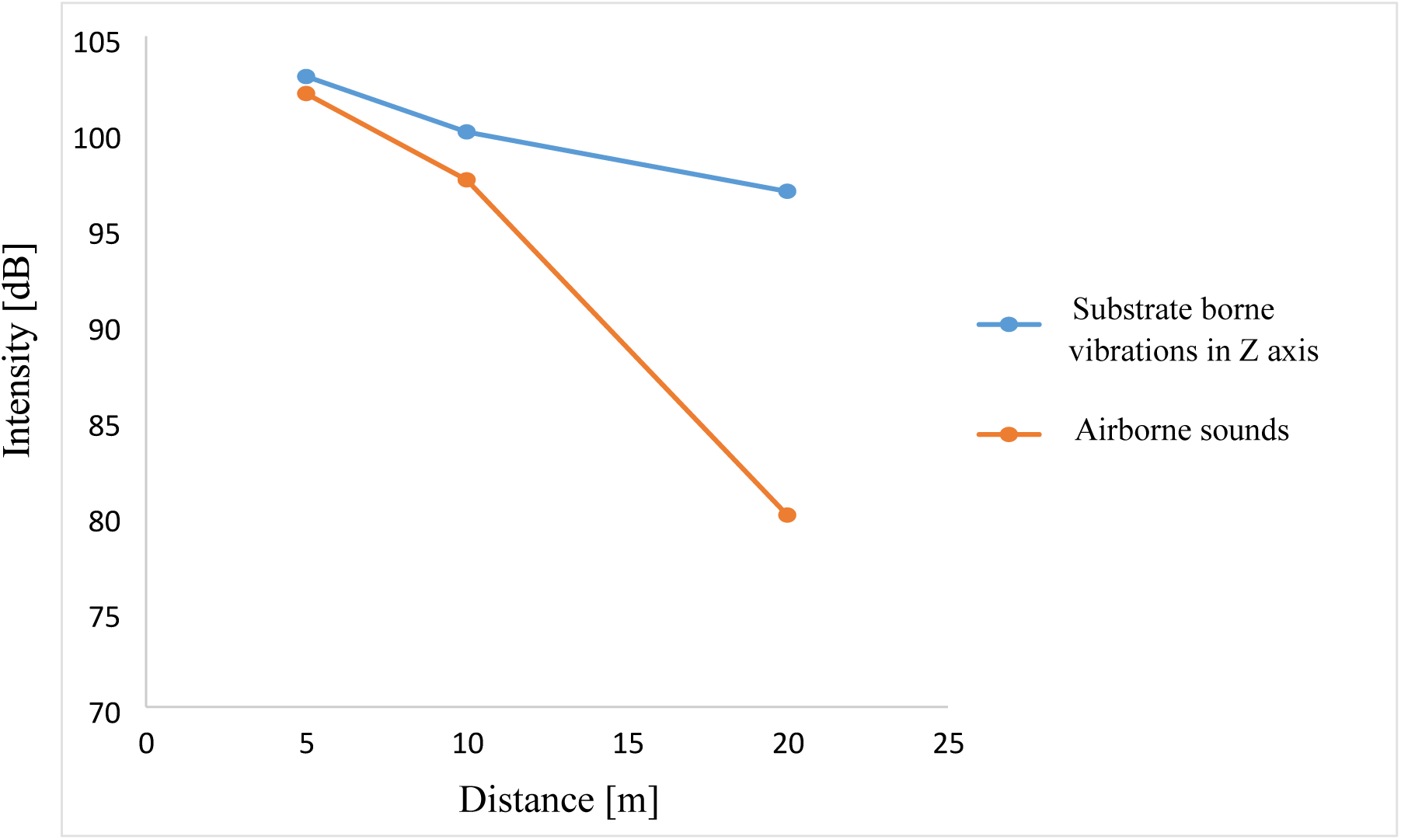
Comparison of substrate borne vibrations (Z axis of all sites; dB re 10 µm/s²) and airborne sound (Düsseldorf A, Düsseldorf B, Dinslaken; dB re 20 µPA =SPL) amplitude over distance in respect to the railway (N trains=679).

### Playback experiments

Individual bees reacted behaviourally to the passage of the train (Chi-square-four-field test: p<0.05; Fig. 4).

**Fig. 4.**
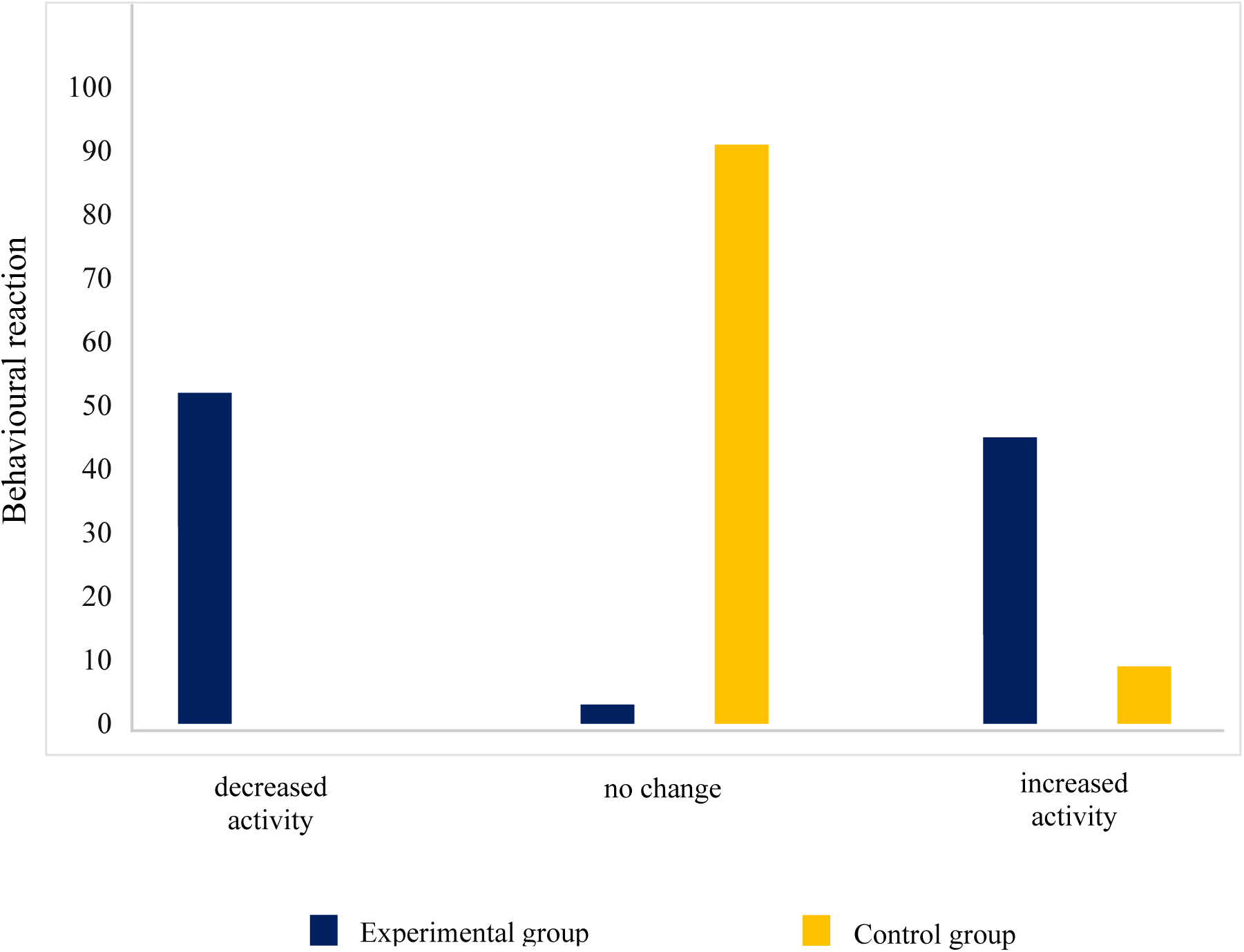
Individual (N=50) behavioural reaction during the experimental and control trials. During the experimental trials (blue bars), bees reacted with decrease or increase of activity; while during the control trials (orange bars), almost no change in behaviour was observed (Chi-square-four-field test: p>0.05).

During the experimental trials, bees showed behavioural reactions as freezing and breaking of their current activity (decreased activity) or more rough chaotic movement (increased activity). In the beginning of the experiment bees tended to react with freezing or almost completely ceased activity (e.g. stop fanning). While the playback continued, bees reacted more with rough chaotic movements (i.e. agitation). In the last part of the experiment, when the colony was more active, the reaction observed was milder. In this case, for some bees the freezing was maintained, others continued their activities or movement with very slow pace, after the initial freezing, or others with accelerated movement. Bees occupied in trophallaxis always ignored the substrate borne vibrations during the whole experiment and occasionally also few bees entertained with other activities ignored the disturbance. Nevertheless, the majority of bees showed a behavioural reaction (∼70%). The clearest effect of the train passage, however, was the lack of buzzing sound right after the passage of the train. Successively, the buzzing sound has slowly increased again.

Both groupś worker population developed normally throughout the season. The worker population was slightly but significantly increased under the experimental conditions in June (U-test: p=0.0159; overall first half season: p=0.0159) and July (U-test: p=0.0317; overall second half season: p>0.05). In the beginning and towards the end of the season there were no significant differences in worker population. No differences between experimental and control group are found in the amount of capped worker brood, worker larvae and worker eggs (U test; p>0.05). No differences are also found in the amount of drone adult population and capped drone brood (U test; p>0.05). Only in May, the amount of drone open brood is significantly different with more open brood in the experimental group (U test; p=0.0397); however, the overall drone open brood is not significantly different between the experimental and the control group (U test; p>0.05). Moreover, no differences are found in the amount of pollen, honey production and total weight (U-test; p>0.05). Just the number of pollen collecting foragers was slightly, but significantly higher in the experimental group in September (U-test: p=0.0397). No differences between the groups were also found in the number of fallen varroa mites after a treatment with evaporated oxalic acid against *Varroa destructor* in September (U-test; p>0.05). No significant differences of the brood index, compensation index and termination index exist between experimental and control group (U-test; p>0.05).

## DISCUSSION

The results show that trains constitute a source of vibrational noise pollution up to at least 20 m from railway tracks. As reported in previous studies on vibrations generated by trains (Ono&Yamada 1989; Heckl et al. 1996), majority of the energy is concentrated in the frequencies below 1000 Hz, which is also the range in which bees are more sensitive and which is exploited for vibrational communication (see review Kirchner et al. 2022). It is already known that bees react to vibrational signals with the so-called freezing response (Michelsen et al 1986a). The freezing response stop the bees for the duration of the signal. Therefore, vibrations at intensities above the threshold of freezing response may constitute a danger for the bees. It could impede or slow down their activities (cleaning, forager recruitment, food collection, conversion of nectar into honey, grooming, sleeping, queen competition, egg laying etc.), which are essential for the colony survival. Moreover, our results show that the most vibrated axis is the one to which honey bees are more sensitive (Z axis coincides with dorsoventral axis in respect to the bee leg; Kilpinen&Storm 1997). The intensity of the noise on the hive seems to vary with the type of ground, the presence of obstacles, the level, the speed and the weight of the trains. In fact, the source characteristics constitute the initial signal intensity but the material, the geometry and/or the size of the medium influence the type of wave transmitted (longitudinal, transverse, bending, Rayleigh), the travelling distance, the attenuation and the spreading of frequencies (Elias&Mason 2014; Hill 2008). In our results we observe that the ground vibrations are much lower than the vibrations on the base of the hive. This suggests that the vibrations are amplified by the hive (Rayleigh 1916); however, without excluding a further potential impact of airborne sounds. On this regard, it must be taken into account that the substrate borne vibrations travel with an attenuation of 3 dB/doubling distance plus a small excess attenuation, diversely from airborne sounds, which travel with an attenuation of 6 dB/doubling distance plus some variable linear excess attenuation and lose energy in contact with boundaries (Caldwell 2014). Therefore, the impact played by substrate borne vibrations with distance can be expected to be higher in respect to airborne sounds.

At the individual level, bees reacted significantly to the vibrations (coherent to the freezing response reported by Rohrsitz&Kilpinen 1997 and Michelsen et al. 1986a). In the first part of the experiment, the behavioural reaction was much more evident in respect to the second part of the experiment, after the pause, when bees were more active. The higher activity of the bees may have influenced the experimenteŕs evaluation, which may have had more difficulties to discriminate a difference; however, there may be other explanations. Firstly, the bees could have habituated to the vibrational disturbance despite the pause (there is evidence of habituation to vibrations in other arthropods; Acheampong&Mitchell 1997; Bullock et al 2009); secondly, the higher buzzing of the bees could have masked the vibrational disturbance or, further, the disturbance could not simply physically be detected due to the shorter contact between the legs and the comb, when bees were moving around. In view of what reported by Terenzi et al. (2020), Farina (2023) but also in view of the experiments by Frings&Little (1957), Little (1962), Stefanec et al. (2021), another third hypothesis could support that the bees learnt to recognize the disturbance of these specific frequencies as something extraneous and learnt to cope with it. They may have only initially or partially been affected but been able overall to continue their activities. Nevertheless, the absence of buzzing immediately after the train passage constitutes a clear reaction to the disturbance (like to what reported by Bencsick et al. 2024) and confirms the detection of the substrate borne vibrations by the bees.

The playback experiment, however, does not show significant effects of vibrational noise at the population level. The experimental colonies, vibrated as if they were placed at 5 m distance from the railways, preserved a good performance, fitness and survival as much as the control group, despite the presence of substrate borne vibrations, both at 1 m/s² and 3 m/s². Indeed, the adult and brood parameters as the monthly and overall amount of pollen and honey are not significantly different. Nevertheless, it is noteworthy that during the experiment at 1 m/s² the worker population in June (1^st^ June-16^th^ July) is significantly higher in the experimental group in respect with the controls. This situation is maintained also in the first part of the experiment at 3 m/s² (17^th^ July-31^st^ July), but afterwards the two groups do not present worker population differences anymore and the overall population is not significantly different between the two groups. With the current information, it is not possible to clarify this specific result, but few hypotheses may be attempted. Firstly, it is conceivable that the bees were longer confined inside the hive, maybe due to lower locomotion or greater life expectation, and therefore more of them has been counted. Secondly, it could also be that the substrate borne vibrational disturbance limits their activities, therefore requiring the rearing of more bees (June, July), until an adjustment is found (August). Between August and September, both the experimental and the control worker population slightly drop, but the pollen collecting foragers are significantly more in the experimental group. This possibly suggests the need for a greater labour force before the end of the season, when the colony prepares for the winter cluster. Gontarski (1957) states that prepupae are sensitive to substrate borne vibrations; however, our results did not show alterations in the brood development. The brood index, compensation index and termination index were the same between experimental and control groups. The number of adult drones and capped drone brood never differed between the groups. Only in May, a significant difference is present for drone open brood, which may be easily explained with beekeeping intervention. Drones were introduced to keep under control the varroa mites. In September, when the colonies were under evaporated oxalic acid treatment, the substrate borne vibrations did not affect the number of varroa mites.

Our colonies were not in swarming mood, therefore it was not possible to assess any effect on queen development, communication (DVAV, tooting, quacking) and mating. However, considering that queen piping can reach amplitudes up to 6-9 m/s² (Michelsen et al 1986b), it seems unlikely that the signals would be completely masked by the train vibrations at least at the amplitudes used in the current experiments. It would be interesting to see whether queens would prefer to communicate in the pauses between the trains (as other insects do with noise sources; Shieh et al 2012; Sathyan&Couldrige 2021) and whether they would have difficulties to find rivals, when emerged, and to respond when still enclosed in the queen cell.

## GENERAL CONCLUSIONS

This study addresses the problem of vibrational noise pollution in honey bees for the first time. Field work show how railway traffic is effectively a vibrational source up to 20 m of distance from railways, with substrate borne waves travelling not only on the ground but transmitted also to a hive. However, the collected data do not state clear adverse effects on honey bee colonies, suggesting that the bees can cope with the disturbance at these frequencies and amplitude. Nevertheless, this does also not exclude that other parameters, which were not evaluated in this study, could be influenced. Colonies subjected to stronger (e.g. mining, aircraft) and/or continuous substrate borne vibrations could show more evident effects (i.e. long-term freezing or reduced flight; Frings&Little 1957 report 20 minute of quiescence in workers and drones in a hive exposed to continuous low frequencies of sufficient intensity). In addition, further data should be collected in relation to queen and drone development, swarming and mating success. On the other hand, at the individual level bees show effectively a behavioural reaction.

In conclusion, more studies are needed to disentangle the effects of vibrational noise pollution on honey bees, but the current data do not suggest that railway traffic may severely harm honey bee colonies, at least for amplitudes up to 3 m/s². Within a more general view, it may be however advisable to place honey bee colonies far from substrate borne vibrational anthropogenic sources, especially when the amplitude on the hive exceed 3 m/s².

## ACKNOWLEDGMENTS

We thank Prof. Beate Brand Saberi for sharing her curiosity about this topic, triggering our enthusiasm to deepen into it. And we also thank our beekeeper, Susanne Staab, for providing bees and materials.

## AUTHOR CONTRIBUTION

WHK and SC developed the study idea and designed the methodology. SC, MCS, JR collected and analysed the data. WHK, SC, MCS jointly wrote and WHK and SC edited the manuscript. All the authors approved the final version of the manuscript.

## DATA AVAILABILITY

Raw data will be made available from the authors upon reasonable request

## CODING AVAILABILITY

not applicable

## DECLARATIONS

Ethics approval: Not applicable

Consent to participate: Not applicable Consent for publication: Not applicable

Conflict of interest: The authors declare no conflict of interest

